# No Distinct Gut Microbiome Signature in Alzheimer’s Disease: A Reanalysis with Modern Tools

**DOI:** 10.1101/2025.09.09.675220

**Authors:** Gugulethu N. Sakana, Sayumi York, Rosa Alcazar

**Affiliations:** Clovis Community College, Fresno, CA 93730, USA; Notre Dame of Maryland University, Baltimore, MD, 21210

## Abstract

Alzheimer’s Disease (AD) is a neurodegenerative disorder that affects memory, cognition, and behavior. This study reanalyzes a publicly available 16S rRNA sequencing dataset (PRJNA554111) of fecal samples from 43 AD patients and 43 age-and-sex-matched controls to assess differences in microbial composition between the groups. We outlined the relative abundances of major phyla, and identified 137 ASVs across five phyla that were differentially abundant (padj<0.05). We found no distinct pattern of microbial composition distinguishing AD from controls, contrary to the original finding of disease-specific signatures.

## Description

Alzheimer’s Disease (AD) is a chronic neurodegenerative disorder that affects memory, cognition, and behavior (Kozhakhmetov et al., 2024). Globally, AD affects an estimated 22% of individuals over the age of 50 (Gustavsson et al., 2023). Over 7 million Americans are living with AD, and it is the 7th leading cause of death among adults in the U.S. (“2025 Alzheimer’s Disease Facts and Figures,” 2025). As AD progresses, neurons are damaged and die due to accumulation of β-amyloid proteins primarily in brain regions responsible for memory, cognition, and language (Zheng & Wang, 2025). The mechanisms underlying AD pathogenesis remain poorly understood.

Researchers have increasingly explored links between the gut microbiome and neurodegenerative diseases, with a focus on altered microbial composition and/or dysbiosis (Solanki et al., 2023). While a bidirectional connection between the gut and the brain has been described, with microbiota playing a key role, few studies have identified and characterized specific taxa implicated in AD within the gut microbiome (Seo & Holtzman, 2024). Moreover, many existing datasets were originally analyzed with older computational methods and limited reference databases, reducing their relevance today.

In this study we analyzed fecal samples from neurotypical (n=43) and AD patients (n=43) originally sampled by Zhuang et al. in 2018 with updated methodology and databases (Zhuang et al., 2018). Reanalyzing this dataset using ASVs with single-nucleotide precision provided greater resolution than the OTUs used in the original study. Additionally, ASVs are reproducible between studies (Callahan et al., 2017), which facilitates comparison of results across studies.

We found no consistent microbial composition between AD and control samples, contrasting with the original study’s claim that gut microbiota are altered in AD. These findings emphasize the importance of re-evaluating older datasets with newer tools to confirm and clarify the relationship between the gut microbiome and AD.

Analysis of the gut microbiota composition at the phylum level revealed considerable heterogeneity among individual samples (Fig. 1A). There was no distinct trend in microbial relative abundance by age, sex, or between AD and control groups. Samples were primarily dominated by Bacillota and Pseudomonadota. Specifically, 68 samples contained more than 50% Bacillota, and 21 samples contained more than 20% Pseudomonadota. The third most abundant phylum was Bacteroidota, with only 14 samples containing more than 20%. In both AD and control groups, we found at least one sample with an atypical microbial composition. Samples C18 and A37 showed the lowest abundance of Bacillota (14.3% and 10.6% respectively), C18 had the highest abundance of Actinomycetota (83.7%) of all samples, while A37 was primarily composed of Pseudomonadota and Bacteroidota (46.4% and 42.1%) respectively. More than half of all samples (n=54) contained no Verrucomicrobiota. Sample A29 had the highest representation of Verrucomicrobiota (40.1%), second only to Bacillota (44.8%).

**Figure 1).**
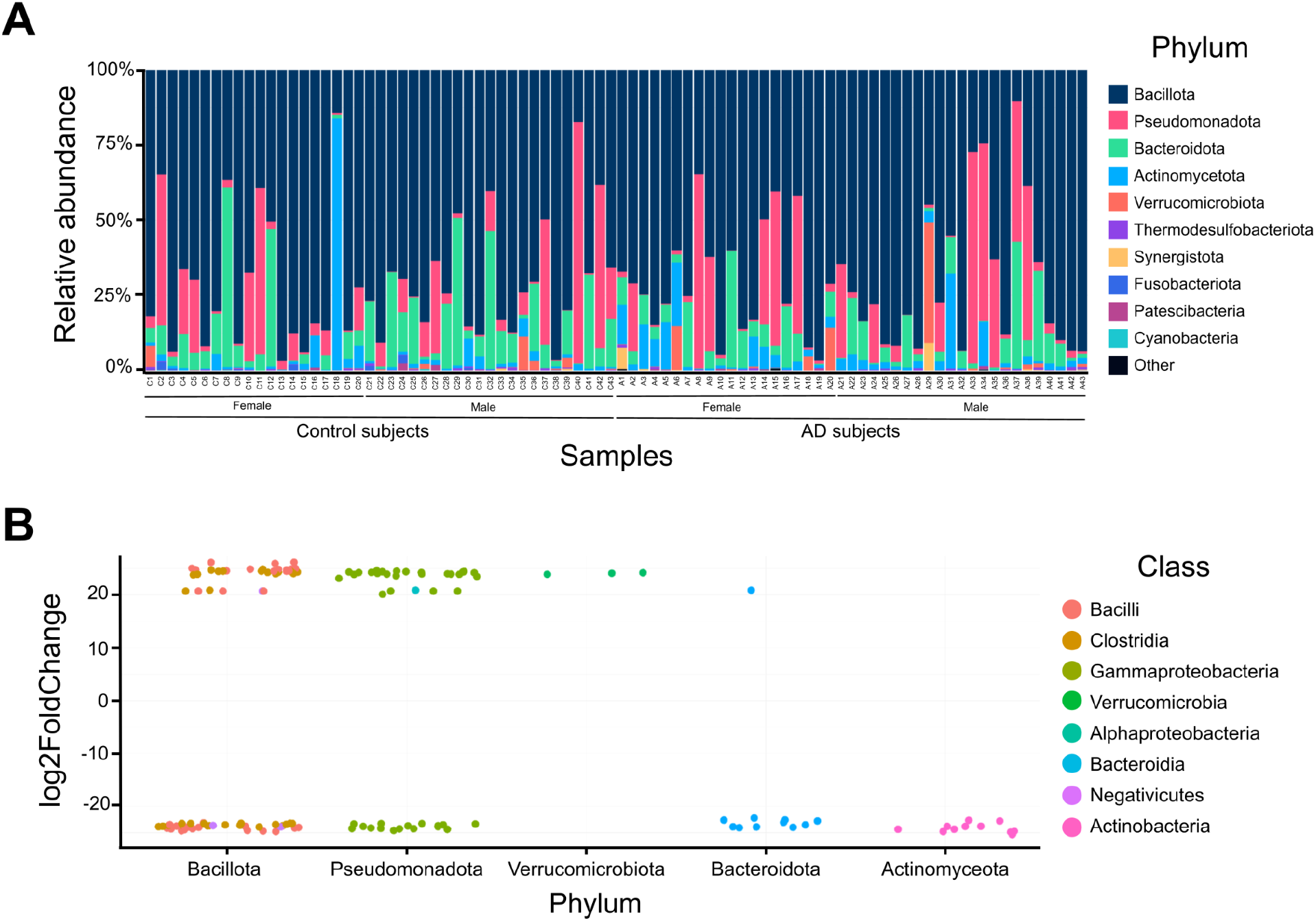
Relative abundance of bacterial phyla per individual and between AD and Control groups. **A**. Barplot shows phylum level composition for each sample, arranged from left to right by increasing age. **B**. Differential abundance plot highlights five phyla with significantly different ASVs (p<0.05) between AD and controls. Positive Log2FoldChange values indicate taxa enriched in AD, while negative values indicate taxa enriched in controls.

We found 137 significantly differentially abundant ASVs (padj<0.05) across five major phyla: Bacillota, Pseudomonadota, Verrucomicrobiota, Bacteroidota, and Actinomycetota. The majority were in Bacillota (70/137) and Pseudomonadota (42/137) (Fig.1b). Within Bacillota, 33 ASVs were more abundant in AD, whereas 37 were more abundant in controls. Within Pseudomonadota, 26 ASVs were more abundant in AD and 16 were more abundant in controls. In both Bacteroidota and Actinomycetota, eleven ASVs each were more abundant in controls than AD. Only one ASV in Bacteroidota was more abundant in AD than in controls. Verrucomicrobiota had the fewest differentially abundant ASVs (3 total), all of which were more abundant in AD than in controls.

The gut microbiome plays an important role in overall human health. The pathogeneses of neurodegenerative diseases like AD remain poorly understood and no curative therapies exist. The present study reanalyzes a dataset originally published by Zhuang et. al (2018) using updated bioinformatics tools and databases to re-evaluate previous findings and enable future comparisons with other studies.

While Zhuang et al. found that microbial composition and diversity were distinct in AD patients compared to healthy age-and-sex-matched controls (Zhuang et al., 2018), we found no consistent compositional pattern relating to age, sex, or disease state. This was reflected in both the lack of commonality within groups and similarities observed across groups. Sample C13 was almost entirely composed of Bacillota, whereas C18 contained the least Bacillota among controls. In the AD group, samples from individuals (A9 and A10) of the same age and condition (67 year-old, AD) showed vastly different abundances of Bacillota and Pseudomonadota. The same was true for individuals of the same sex. Samples C17 and C18 were both from female subjects, but C18 was dominated by Actinomycetota and C17 contained almost none; a similar pattern among male subjects was exemplified by samples A31 and A33. In contrast, samples in AD and control groups were sometimes strikingly similar. A27 and C39 had the lowest abundance of Pseudomonadota, and contained similar levels of Bacillota and Bacteroidota. Samples from individuals of different ages (C7, 62 years; A16, 79 years) also had very similar abundances of Bacillota, Pseudomonadota, and Bacteroidota. Studies have also shown inconsistent observations of relative microbial abundance among patients with other neurodegenerative diseases such as Parkinson’s Disease (PD) and Amyotrophic Lateral Sclerosis (ALS) (Rob et al., 2025).

Zhuang et al. excluded subjects from the healthy control group if they had a family history of neurodegenerative disease or a diagnosis of major chronic illnesses such as gastrointestinal disease or a diagnosed mental illness, but control subjects may have other underlying unknown chronic health conditions (Zhuang et al., 2018). Additionally, Zhuang et al’s study included AD patients diagnosed by experienced neurologists, but of the 43, only 12 underwent positron emission tomography (PET) inspection for deposits of β-amyloid in the brain. Unfortunately, data on which of the AD subjects underwent PET or the severity of their AD diagnosis are not available. These factors may contribute to the absence of a microbial composition pattern clearly distinguishing AD from controls. Notably, other research on the gut microbiome has found the individual to be the largest factor in explaining microbial composition, above age and sex (Guthrie et al., 2022).

We found five major bacterial phyla (Bacillota, Pseudomonadota, Verrucomicrobiota, Bacteroidota, and Actinomycetota) that were differentially abundant in AD patients compared to their healthy age-and-sex-matched controls, whereas differential abundance at the phylum level previously only revealed differences in Actinomycetota and Bacteroidota. Actinomycetota was originally found to be significantly more abundant in AD (Zhuang et al., 2018), but our results showed the opposite, with all differentially abundant Actinomycetota ASVs more abundant in healthy controls.

Systematic reviews have revealed that studies found taxa within Bacillota to be reduced in the gut microbiota of some AD patients (Murray et al., 2022). However, others have found taxa within Bacillota, Bacteroidota, and Pseudomonadota are not significantly different between healthy controls and AD patients (Heravi et al., 2023). Bacteroidota can be both beneficial and opportunistic pathogens in some parts of the gut, as they also produce short-chain fatty acids (SCFAs), but can invade other locations and cause intra-abdominal, skin, respiratory, and brain abscesses (Patrick, 2022; Shin et al., 2024). While an elevation of the abundance of taxa within the phylum Pseudomonadota in the gut has previously been implicated in AD (Kozhakhmetov et al., 2024), our results do not support a correlation between relative abundance of Pseudomonadota and AD; ASVs in the phylum did not show a consistent pattern in abundance between control and AD samples. Analysis at greater levels of taxonomic resolution would be useful in untangling this relationship further. Actinomycetota, a relative minority in the gut, plays a key homeostatic role. In particular, Bifidobacteria help maintain the gut barrier via production of SCFAs like acetate, which help protect the host from pathogenic infection (Binda et al., 2018).

AD, like many neurodegenerative diseases, remains a complicated, unsolved puzzle. Clinical presentations of AD are highly heterogeneous and diagnosis typically involves a multifactorial analysis of the individual (Atri, 2019). The disease has more recently been described as a spectrum ranging from asymptomatic, preclinical stages to mild cognitive impairment and dementia. Our results reflect the complexity of AD development and progression; showing no clear patterns in microbial composition by diagnosis, age, or sex. Our study also demonstrated the importance of re-examining older datasets with updated pipelines, illustrated by our finding that Actinomycetota ASVs were more abundant in controls, contrary to the previous report.

Given the conflicting findings on gut microbial composition between AD and controls in this study and in the broader literature, further work should focus on reanalyzing additional datasets to confirm or refute foundational claims about the link between the gut microbiome and AD.

## Acknowledgements

This work was supported by NIH grant UE5HG013799-01. We thank members of the C-MOOR team, as well as fellow C-MOOR Scholars Gauri Paul, Grace Ekalle, and Madeleine Gerard for their support and advice throughout the process.

## Methods

Data from Zhuang et al. (2018) were downloaded from the NCBI SRA (BioProject PRJNA554111) and processed in DADA2 (Callahan et al., 2016; Zhuang et al., 2018). In brief, sequence quality was evaluated using the *plotQualityProfile* function and processed using the *filterAndTrim* function. Forward and reverse reads were trimmed to a length of 250bp and reads with more than 1 expected error were removed; all other settings remained at default values.

Samples were then dereplicated and forward and reverse reads were merged using the *derepFastq* and *mergePairs* functions. Chimeric sequences were then removed using the *removeBimeraDenovo* function. Taxonomic assignment was performed using the assignTaxonomy function to the level of species using the SILVA 138.2 database (Quast et al., 2013).

We analyzed and visualized the microbiome composition of samples using the R packages phyloseq and ggplot2 respectively (McMurdie & Holmes, 2013; Wickham, 2011). Differential abundance analysis was performed in DESeq2 using the alternative geometrical mean to estimate size factors and a *padj*-value cutoff of 0.05 to identify ASVs that were differentially abundant between the control and AD groups (Love et al., 2014).

